# Prevalence and environmental abundance of the elusive membrane trafficking complex TSET in five cosmopolitan eukaryotic groups

**DOI:** 10.1101/2024.06.15.599112

**Authors:** Mathias Penot-Raquin, Mandeep Sivia, Morenikedji Fafoumi, Raegan Larson, Richard G. Dorrell, Joel B. Dacks

## Abstract

Eukaryotic cell biology is largely understood from paradigms established on few model organisms, largely from the animal and fungi (opisthokonts) and to a lesser extent plants. These organisms, however, constitute only a small proportion of eukaryotic diversity, and the principles of their cell biology may not be universal to other, understudied but globally impactful, organisms. Intriguingly, there are cellular components that are present in diverse eukaryotes, but are not in the animals and fungi on which the best developed models of cell biology are derived. Consequently, these components are not included in the generally adopted frameworks of cellular function that are meant to explain eukaryotic biology. The membrane complex TSET is the best studied such example, well established to play a role in cell division and endocytosis in plants. It is found across eukaryotes, but is highly reduced in opisthokonts. Its general prevalence, abundance, and relevance in eukaryotic cellular activity is unclear. Here we show that TSET is encoded in genomes of five cosmopolitan and critical groups of primarily photosynthetic eukaryotes (green algae, red algae, stramenopiles, haptophytes and cryptophytes), with particular prevalence in the green algae and some stramenopile groups. A meta-analysis of published gene expression data from the model diatom *Phaeodactylum tricornutum* shows that this complex is coregulated with components of the endomembrane trafficking machinery. Moreover, meta-transcriptomic data from *Tara* Oceans reveals that TSET genes are both present and expressed by diatoms in the wild. These data suggest that TSET may be playing an important and underrecognized role in cellular activities within marine ecosystems. More broadly, the results support the idea that use of systems-level data for non-model organisms can illuminate our understanding of core principles of eukaryotic cell function, and may reveal important and under-appreciated players that deserve to be integrated into the pervasive models of cellular capacity.

## Introduction

The vast majority of eukaryotic diversity is found in microscopic, i.e. protist, lineages [1]. Over the past two decades, next-generation sequencing of genomes and transcriptomes has transformed our view of the evolutionary and cellular diversity of life [1,2]. While the exact taxonomic structure of eukaryotic diversity remains continually challenged, e.g. by the recent identification of new phyla and even kingdoms of microbial eukaryotes [3,4], phylogenomic analysis of protist genomes and transcriptomes broadly resolves all eukaryotes into between five and eight “supergroups” [1].

There is a disconnect, however, between our understanding of eukaryotic diversity and our understanding of eukaryotic cell biology. Most model organisms used to study eukaryotic cell biology are confined to one or perhaps two, primarily multicellular groups; the opisthokonts which encompasses the vast majority of model organisms such as humans, mice, rats, flies, worms, and yeasts; and to a much lesser extent embryophytes, dominated by a few model plants. The other major eukaryotic groups (i.e. Amoebozoa, CRuMs, Provora, Hemimastigophora, Metamonada, Discoba, Archaeplastida, Haptista, and TSAR ) are populated by protist and algal groups whose biology has been comparatively understudied, and yet are of massive medical, biotechnological, agricultural, and environmental relevance (reviewed in [1]). For questions that address cellular function, it is imperative that our cellular understanding accurately encompasses the biology in these organisms, and not just the opisthokont and embryophyte models. While this certainly true for specific questions of particular organisms, this diverse assemblage of unicellular eukaryotes has the potential to transform, and challenge, our view of how the majority of conventional eukaryotic cells function [5].

One reason for this historical disconnect has been the limited amenability of many eukaryotic micro-organisms for laboratory investigation, notwithstanding the outstanding contribution from studies of a few key model parasitic organisms. Some protists inhabit ecosystems that are difficult to access (e.g., deep-sea sediments, the open ocean, polar ice-caps), or may be difficult to establish into stable laboratory culture [6]. This led on to a lack of tools for investigation or manipulation. However, technical advances are erasing these barriers. Environmental sequencing (i.e., meta-genomics) has cast new light into the diversity and function of the most abundant eukaryotic micro-organisms in the wild, which may elude characterisation from classical cultivation. This approach also expanded our understanding of previously cultured groups (e.g., dictyochophytes within the stramenopiles) that are nonetheless far more abundant in the wild than in culture collections [7]. It has also allowed the identification of novel eukaryotic groups (e.g., rappemonads within the haptista) that had not been placed into culture when they were first identified from environmental (typically 18S rDNA) sequencing [8]. Environmental sequencing has not only expanded our view of protist genetic diversity, but has also allowed us to identify cultivation roadblocks that, when solved, can give us a more complete view of eukaryotic cell organisation [6,9,10]. Once representatives have been brought into the lab, systems-level data is being collected in the form of gene expression studies, and proteomics. Moreover, there has been a concerted effort for development of genetic and cell biological tools to expand and rebalance the diversity of culturable models for eukaryotic cell biology [11]. This has led to discoveries of widespread cellular phenomena, a “cultural renaissance” [12].

One intriguing set of cellular factors are the so called “jötnarlogs” [13]. These are defined as, “a homolog with the evolutionary pattern of being ancient but hidden from view, as with the dark or walled-off ancient world of Jotunheim in Norse mythology. Specifically, genes are jötnarlogs if found in sufficiently diverse eukaryotic taxonomic supergroups to infer a common origin concurrent with or pre-dating the LECA *(Last Eukaryotic Common Ancestor)*, but were hidden from previous cell biological investigation due to loss or divergence in yeast and animal model systems” [13]. Within just the membrane-trafficking system there are, at least, 12 examples [13,14]. The existence of such a category of genes is established but raises urgent follow on questions. How taxonomically prevalent are such genes: are they rare curiosities or underappreciated common cellular components? Are jötnarlogs found in important microbial players; drivers of environmental systems and medically relevant organisms? And is there evidence that the components are actively playing roles in such organisms or are they merely minor factors, such that the existing cell biological models based on opisthokont biology still capture the overall picture of cellular function?

One of the best studied jötnarlogs is the membrane trafficking complex, TSET. Originally identified in the model plant *Arabidopsis thaliana* [15], TSET (or the TPC as it is called in plants) was first considered to be embryophyte-specific, but was subsequently detected in the distantly-related amoebozoan *Dictyostelium discoideum* [16]. It is an adaptin-related complex, composed of six subunits (named TSPOON, TCUP, TPLATE, TSAUCER, TTRAY1 and TTRAY2) which is homologous but distinct from the five conventional adaptins and the COPI complexes which all serve as cargo-recognition machinery in vesicle formation at various organelles of the endomembrane system [17]. TSET similarly serves important functions in cell trafficking and turnover. It is best characterized in plants where it plays roles in phragmoplast deposition and cell division [18], actin-mediated autophagy [19], as well as clathrin-mediated endocytosis [15,20]. This latter function at least is shared with *D. discoideum*, albeit whether clathrin is also involved in TSET mediated endocytosis in this specific case is unclear [16]. In plants, TSET may be as important for this process as the adaptor complex AP2 [15], which is the major mediator of this process in animal and fungal cells. Thus, TSET may well underlie endomembrane system function, endocytosis or otherwise, in a broad range of eukaryotes, but is not generally integrated in models of cell surface uptake. Comparative genomics established that TSET is pan-eukaryotic [16]. However, opisthokonts retain only the TCUP subunit (FCHO in animals and Syp1 in yeast, which nonetheless has a role as an accessory protein in Clathrin-mediated endocytosis [21]. Notably, in the initial study reporting pan-eukaryotic distribution of TSET [16], the patterns of retention were relatively consistent within most taxonomic groups assessed (i.e. nearly all subunits either detected or not).

However, this was not the case for the major groups of algae, where the pattern of detected TSET subunits was far more sporadic. The green algae, red algae, cryptophytes haptophytes, and stramenopiles are of critical environmental relevance but contrasting evolutionary and functional diversity. Those lineages are distantly related to animals, fungi, plants, and even one another in the eukaryotic tree (Fig S1). They are mainly composed of photosynthetic organisms with algal lifestyles in marine and freshwater environments, and thus play major role in the primary production of aquatic ecosystems. Diatoms, within the stramenopiles, alone account for 30-40% of global primary production [22]. Moreover, several lineages are mixotrophic (haptophytes) or fully heterotrophic, such as the plant pathogens oomycetes (stramenopiles), the human gut commensal *Blastocystis* or the phagotrophic genus *Goniomonas* (cryptophytes). Taken together, these five groups also represent > 99% of the total algal 18S and plastid 16S rDNA abundance found in the *Tara* Ocean campaign [6,7]. Many marine algae exhibit rapid turnover and short generation times in blooming cycles that may enable them to capitalise on the transient nutrient availabilities in most ocean environments [23]. Similarly, most eukaryotic algae are often characterised by a reliance on phagotrophic strategies alongside photosynthesis (i.e., photo-mixotrophy) and endocytotic mechanisms for nutrient uptake [24]. Thus, the same physiological processes in which TSET is implicated in plants are naturally relevant to the ecophysiology of marine microbes.

In the original TSET pan-eukaryotic survey [16], two of the green algae were found to possess the entire complex, while the others had a more sporadic distribution, as did the single representatives of the glaucophytes, haptophytes, and cryptophytes assessed. The single red algal representive appeared to encode no TSET subunits. Most puzzlingly, the four stramenopile genomes sampled at the time of TSET discovery (i.e. the diatom *Thalassiosira pseudonana*, kelp *Ectocarpus siliculosus*, eustigmatophyte *Nannochloropsis gaditana*, and oomycete *Phyophthora sojae*) revealed an incomplete TSET complex in each lineage, with all stramenopiles possessing fewer than three of the six documented subunits and lacking TPLATE. However, a recent study identified a complete TSET complex in the mixotrophic green algae Cymbomonas [25], and a complete TSET has recently been documented in the hyper-abundant gut microbe *Blastocystis* and its relative *Proteromonas lacertae* [26], both members of the stramenopiles. These data suggest that TSET may be a larger player in cellular processes in algae, and its taxonomic prevalence, putative function, and relative abundance beg to be explored.

Since 2014, the genomic, transcriptomic, and environmental sequencing data available for microbial eukaryotes, including marine algae has massively expanded. Therefore, here we revisit the question of TSET distribution and prevalence in these algal and related group, using an expanded library of over 700 stramenopile, haptophyte, green algae, red algae and cryptophyte algal genomes, transcriptomes and meta-assembled genomes. We apply an iterative homologue retrieval (AMOEBAE) approach [27], and phylogenetic validation to assess the TSET diversity across these lineages. We further use transcriptome data from the model diatom species *Phaeodactylum tricornutum*, and diatom meta-transcriptomic data from the *Tara* Ocean expedition, to assign preliminary functions to the TSET complex, at least in a model diatom. Our work shows that this membrane trafficking complex is widespread and complete in several lineages of eukaryotic algae and their close heterotrophic relatives. We further show that this complex is expressed by wild diatoms in the open ocean, and appears to have a function in a central process of cell biology in a model diatom in the lab. Our study addresses the lingering questions of jotnarlog relevance to cell biological models (at least for TSET) and emphasises the importance of considering cultivated and uncultured microbial diversity of critical ecological importance for exploring fundamental eukaryotic cell processes across the eukaryotic Tree of Life.

## Methods

### Reference library construction

First, a composite reference library of 266 stramenopile, 44 haptophyte, 281 green algal, 42 red algal and 34 cryptophyte genomes and transcriptomes (Dataset S1) was compiled, extending from previous studies [28,29]. Broadly, this consisted of publicly accessible genomes, accessed from JGI PhycoCosm (07/22), decontaminated MMETSP and OneKp transcriptome libraries for each group, plus eighteen further chrysophyte and seven further diatom (both stramenopile) transcriptomes independently sequenced in other studies. Libraries were hierarchically organised according to the results of recent multigene phylogenies of each lineage [30–32] and were divided into groups of broadly equivalent rank to orders. For example, stramenopiles were divided into Oomycetes, Labyrinthulomycetes, Pelagophytes, Pinguiophytes, Dictyochophytes, Phaeophytes and Xanthophytes (PX), Raphidophytes, Diatoms, Eustigmatophytes, Synurophytes and Chrysophytes (SC), Bolidophytes, Bicosoecids, and *Blastocystis*; and haptophytes were divided into Pavlovophytes, Phaeocystales, Prymnesiales, Isochrysidales, and Coccolithales-Syracosphaerales-Zygodiscales (CSZ). The full taxonomically sorted library is available for user exploration at https://osf.io/ykxes/.

### Homologs detection

Searches for putative TSET subunit homologs were performed using the AMOEBAE workflow. Protein sequences of confirmed TSET components (Dataset S2) were collected from Hirst et al. [16] and aligned on Seaview using ClustalO (default parameters) [33,34]. These alignments were then used as queries in AMOEBAE, a semi-automatic reverse blast search tool. Several reference genomes were used for the AMOEBAE searches: *Arabidopsis thaliana* (GCA_000001735), *Dictyostelium discoideum* (GCF_000004695), *Phaeodactylum tricornutum* (GCA_000150955), *Blastocystis hominis* (GCA_000151665) and *Emiliana huxleyi* (GCA_000372725). The same queries, reference genomes and Ref_seqs_1_manual_predictions.csv files were used for independant searches in stramenopiles, haptophytes, red algae, green algae and cryptophytes datasets. For Stramenopiles and Haptophytes, searches were performed using a dataset composed of both available NCBI reference genomes (accessed 10/10/2023) and the custom reference dataset supplied above (Dataset S1). For each AMOEBAE searches, all four AP1 subunits sequences from the fungi *Rhizophagus irregularis* were used as positive controls, this complex being present in every eukaryote examined thus far [27]. Organisms in which only less than two AP1 subunits were detected were considered as not complete enough and thus not retained for further analyses.

For each TSET subunit found in stramenopiles and haptophytes, sequences of the putative homologs were retrieved and aligned with the query sequences and their AP/COP paralogues (outgroup) in Seaview using ClustalO (default paramters) [33,34]. Alignments were trimmed in Seaview using Gblocks (options for a less stringent selection) [33,35], and phylogenetic trees were computed in Seaview using PhyML with default parameters (LG model, aLRT branch support). Putative homolog sequences that did not group with the reference sequences were considered as false positives and removed using the Taxus software. Successive rounds of selection were performed to obtain a topology with a clear monophyly of the TSET subunit sequence. Complete homologs lists, alignments, and curated tree topologies are provided in Dataset S3.

### Phaeodactylum co-regulation analysis

To identify genes co-expressed with TSET subunits in the *P. tricornutum* genome, a meta-dataset of previously published gene expression data was constructed following previous studies [28,36]. Broadly, this consisted of two normalised meta-studies of RNAseq (PhaeoNet) [37] and microarray data (DiatomPortal) [38], with relative transcript abundances or fold-changes (respectively) transformed into ranked values, where 100 corresponded to the most highly expressed gene in the dataset, and 0 the lowest. Genes not evaluated in each study were marked as N/A to avoid biasing the correlation values obtained. Pearson correlations of ranked gene expression abundances (i.e., Spearman correlations) were then calculated for all genes in the version 3 annotation of the *P. tricornutum* genome [39] and six genes retrieved to encode potential TSET subunits: Phatr3_J43047 (TCUP1), Phatr3_J43761 (TCUP2), Phatr3_J54511 (TPLATE), Phatr3_J54718 (TSPOON1), Phatr3_J14536 (TSPOON2), and Phatr3_J46356 (TTRAY). An average correlation value was calculated for all six subunits, and for a “core” set of TCUP1, TPLATE, TSPOON1 and TTRAY, as TCUP2 and TSPOON2 were found to have limited transcriptional coregulation with other TSET subunits.

*Phaeodactylum* genes were annotated following previous studies for identified epigenetic marks (histone acetylation and methylation, DNA methylation, and polycomb group presence), predicted function (GO, PFAM, KEGG), and predicted localisation of the encoded protein (ASAFind, HECTAR, MitoFates, WolfPSort) [37]. Complete tabulated coregulation coefficients and predicted protein annotations are provided in Dataset S4. The KEGG Mapper Pathway (accessed 26/10/2023) was fed with all K numbers listed in the Dataset S4 in order to determine the cellular processes correlated with the expression of the “core” TSET genes, with a correlation coefficient threshold > 0.5. The pathway composition obtained with all the gene from the version 3 of the *P. tricornutum* genome was used as a reference to calculate the completeness of each pathway after the correlation filter, i.e. the percentage of genes from a specific pathway which are coregulated with the “core” TSET genes.

### Tara Oceans analysis

For each TSET subunit, diatom homologues were aligned in Seaview using ClustalO [33,34]. Alignments were converted in stockholm format using the online Bugaco converter (http://sequenceconversion.bugaco.com/converter/biology/sequences/fasta-2line_to_stockholm.php), and then converted into a HMM file using the hmmbuild function of HMMER 3.3.2.

Putative homologous transcripts were retrieved from the Tara Oceans metatranscriptomic dataset thanks to the online Ocean Gene Atlas tool (https://taraoceans.mio.osupytheas.fr/ocean-gene-atlas/), using the following options: MATOUv1+T dataset, 1×10^-10^ expect threshold, abundances as percent of total reads, and the custom HMM file as the query. The putative homolog abundances were downloaded, as well as the associated environmental parameters. Only sequences previously annotated as diatoms (Bacillariophyta) were used for analyses. We believe this preliminary taxonomic annotation is sufficient to classify diatom TSET sequences due to the absence of evident horizontal transfers of its subunits within cultured species. Transcript abundances were normalised for each station, depth and size fraction using the total transcript abundance for diatoms in the corresponding sample, in order to prevent biases due to the ecological preferences of these organisms [40]. Analyses were conducted using R version 4.2.1 and the ggplot2, corrplot, factoextra, FactoMineR, VIM and missMDA packages.

For TSAUCER, only one diatom homologue was available. As the blastp search on OGA (same parameters as described above) did not retrieve any homologous transcripts, a hmmer search was performed using the TSAUCER reference query sequences alignment (see Homologs detection), with no success.

## Results

### Widespread occurrence of TSET across marine algal groups

To address the question of whether TSET in marine algae is a rare curiosity or a more prevalent fixture, we assembled an extensive dataset that included genomes, transcriptomes, and environmental metagenomic assembled genomes. This approach is biased toward broad taxonomic inclusion, intentionally designed for positive detection of orthologues in as many lineages as possible. However, it includes datasets with potential low coverage, and thus risks false negative errors. This prevalence assessment is likely an underestimate of the true state.

We began with an assessment of TSET in green algae. Previous studies had reported that the full TSET complex was encoded in landplants, *Chlamydomonas, Volvox*, and *Cymbomonas*, but sporadically assessed TSET prevalence elsewhere [16,25]. Here we were able to identify each of the six TSET subunits in 5 of the 6 groups, with only the Nephroselmids lacking TCUP and TTRAY2 (Figure 1A, Dataset S5). There were 17 instances of the complete TSET complex being identified within a given organismal dataset, including new reports from *Chlorella, Chlor, Coccomyxa*, three *Nannocloris*, three UTC transcriptomes and three datasets from the Chlorodendrophytes-Pedinophytes, as well as *Chlamydomonas, Cymbomonas*, and *Volvox* as previously reported. The Mamiellophytes most frequently lacked TCUP and TTRAY2, with TCUP being the most frequently undetected subunit across all datasets. Overall, this suggests that the ancestor of green algae likely possessed a complete TSET complex and indeed, that TSET is likely a substantial component of modern green algal trafficking systems, especially in the UTC clade, which can represent dominant green algae particularly in coastal regions [41].

**Figure 1.**
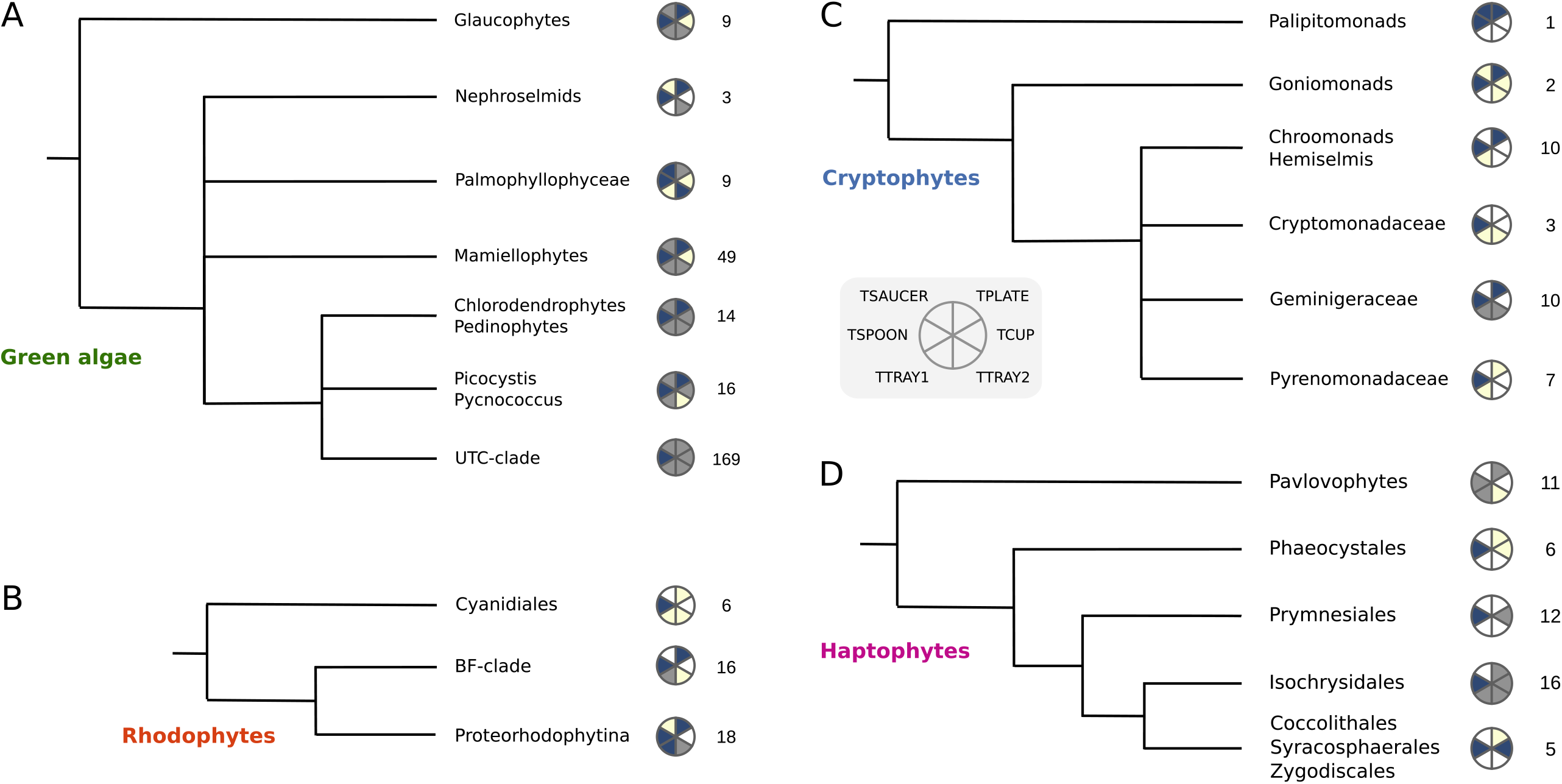
Prevalence of TSET in major marine algal groups. Illustration of TSET subunit detection in the subgroups of each of four algal groups. Light yellow sectors indicate that a unique ortholog was detected. Grey sectors indicate that orthologues were detected in less than 50% of the datasets. Blue sectors indicate that orthologues were detected in ≥50% of the datasets examined. The number of genomes/transcriptomes inspected is indicated on the right side of the corresponding group. A: Green algae and glaucophytes. B: Red Algae. C: Cryptophytes: D: Haptophytes.

Related to the green algae and embryophytes within the supergroup Archaeplastida are the rhodophytes and glaucophytes. Despite the comparatively small sampling of glaucophyte genomic datasets (Fig 1A, Table S5), we were able to detect each of the TSET subunits, albeit with TCUP only found in a single case and never with a complete complex in a given database. In the red algae, we observed an overall paucity of TSET subunits (Fig 1B, Dataset S5). A similar paucity of TSET was observed in cryptophytes (Fig 1C, Dataset S5). Each of the subunits was identified in cryptophytes, but in no case did we see an example of the entire complex in a given organism. Most notably, this trend continued in the haptophytes. A previous analysis identified TCUP subunits in *Emiliania huxleyi* and two related coccolithophores [42]. This was confirmed here, along with individual examples of each other subunits except TSAUCER. However, we were unable to find a complete TSET (Figure 1D, Dataset S5), in any of the 53 genomes and transcriptomes inspected including *E. huxleyi*, for which multiple complete genomes and transcriptome assemblies exist [43]. While the potential for false negatives is high in our analysis, overall these data suggest the complete TSET complex seems unlikely to be a prominent factor in haptophyte cellular trafficking as a complete complex, as it does in plants. That said, a TSPOON subunit was identified in nearly all of our sampled haptophyte databases (Dataset S5). In opisthokont systems, TSET has been reduced to a single remnant TCUP subunit, that acts as a clathrin accessory subunit [21]. The near complete retention of TSPOON in our haptophyte datasets leads to a hypothesis that it could be acting as a monomeric protein, something that would require experimental testing once a model cell biological system has been better developed for haptophytes.

Finally, we examined stramenopiles using both the universal approach as well as an approach using stramenopile queries identified in the first round of searches as queries and reverse BLAST databases in a subsequent round to increase sensitivity. The searches were done into 336 stramenopile genomes and transcriptomes (Dataset S1), grouped into 13 taxonomic lineages, and putative homologues were validated by phylogenetic analysis (Fig 2, datasetS5). We identified all six components of the TSET complex in four stramenopile groups; dictyochophytes, diatoms, the photosynthetic “PX” clade (Phaeophytes and Xanthophytes) and in the gut-denizen, Blastocystis. This latter result is a confirmation of a recently published observation, along with the presence of TSET in the relative of Blastocystis, *Proteromonas lacertae* [26]. However, it is notable since this is a rare case where all six TSET subunits are found in the same genome. In five other stramenopile groups we were able to identify five of the six subunits, and sporadic presence of the complex subunits in all other cases. Individual phylogenies of the TSET subunits broadly recovered the same branching relationships as the species topology, suggesting largely vertical inheritance as opposed to horizontal transfer between different stramenopiles (Fig. S2). Nonetheless, we did observe unexpected patterns of detection of subunits across our dataset and with the different subunits. Leaving aside the differences between datasets from isolates or species of closely related organisms, as likely due to dataset incompleteness, we did note that while TPLATE, and TSPOON were relatively frequently identified in our datasets, the other components were more infrequently found. TSAUCER was particularly rare and with a puzzling distribution. In several cases, a single representative retained a validated homologue, but this was not found in the other datasets of closely related organisms. In the case of Dictyophytes, using the single identified dictyophyte TSAUCER protein did identify two additional, suggesting that this may be explained by rapid sequence evolution that causes detection failure. Nonetheless, use of HMMs made from stramenopile orthologues only and using Blastocystis as the reverse BLAST database did not substantially change the observed pattern of retention (data not shown). Considering the phylogenetic relationships of different stramenopile groups, and the inherent emphasis on taxon sampling, rather than completeness of the datasets, these data are most consistent with presence of a complete TSET in the stramenopile common ancestor.

**Figure 2.**
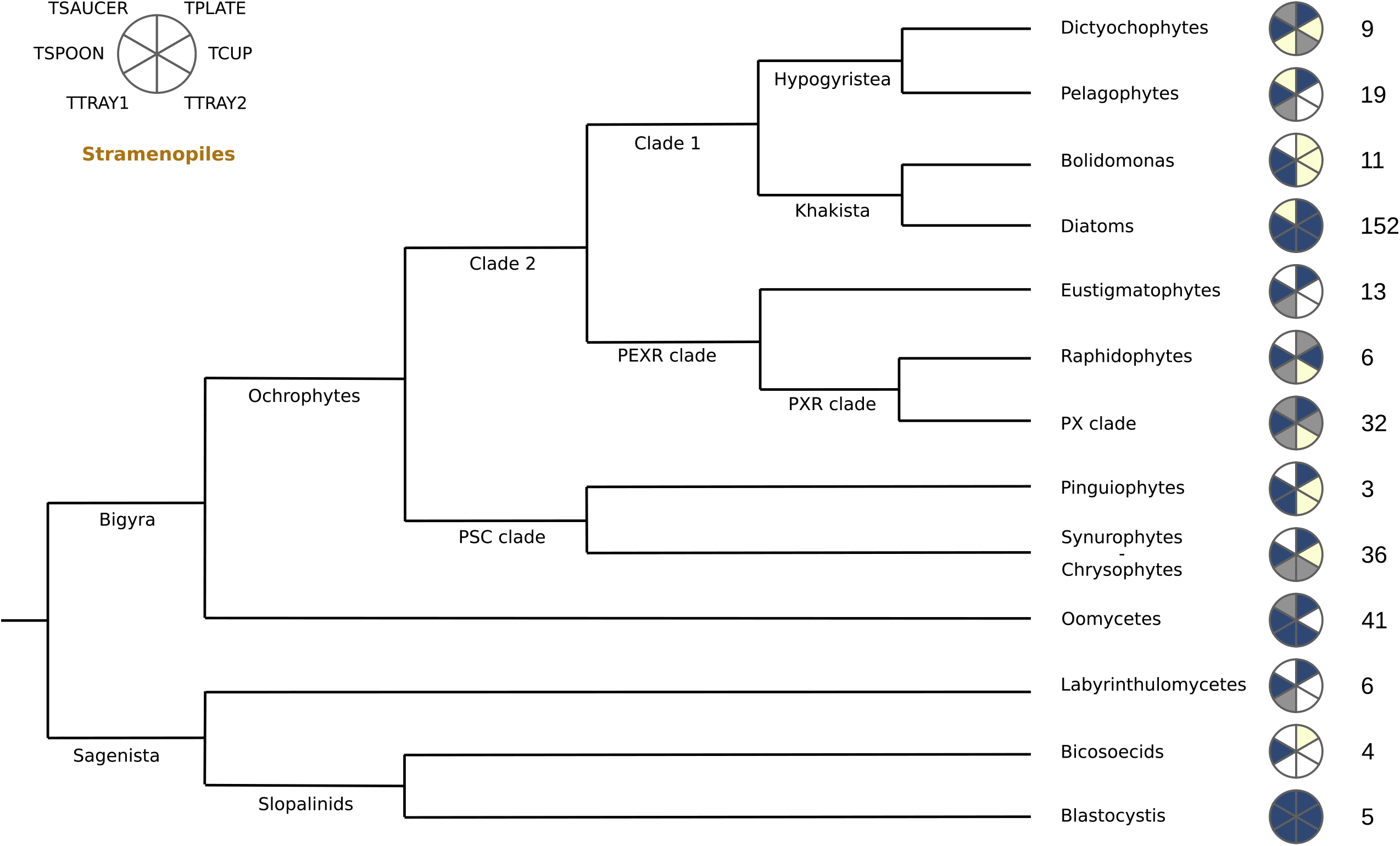
Distribution of TSET across stramenopiles. Light yellow sectors indicate that a unique ortholog was detected. Grey sectors indicate that orthologues were detected in less than 50% of the datasets. Blue sectors indicate that orthologues were detected in ≥50% of the datasets examined. The number of genomes/transcriptomes inspected is indicated on the right side of the corresponding group.

### Coregulation analysis suggests implications of the Phaeodactylum TSET in endomembrane trafficking

The comparative genomic data that we obtained is consistent with TSET having a potential role in membrane-trafficking in stramenopiles, including for medically important stramenopiles such as Blastocystis, agriculturally important organisms such as oomycetes, and environmentally important algae. And yet, the sporadic distribution (and the failure to identify TSAUCER in most taxa) also raises questions about functional relevance. Because of the major role that diatoms play in aquatic environments, we were particularly interested in further investigating this group of algae and whether TSET may be playing similar roles in secretion and endocytosis in diatoms as it does in plants.

To investigate this question, we took advantage of the rich set of systems level data from the model diatom species, *P. tricornutum*. A high-quality and complete genome annotation is available for this organism, as well as extensive gene expression data, considering the epigenome, RNAseq, and microarrays performed under multiple conditions [37,38]. Using a previously benchmarked protocol [36], we identified genes across the version 3 annotation of the *P. tricornutum* genome that show strong positive co-expression trends with six annotated TSET subunits. Considering an average calculated Spearman correlation value of > 0.5 for four central subunits (TCUP1, TPLATE, TSPOON1 and TTRAY) as evidence of positive coregulation, we identified the most strongly positively coregulated genes and functions [36]. The complete data are available as a pivot table (Dataset S4).

Across the entire *Phaeodactylum* genome, we noted coregulation trends that may provide insights into TSET function. The most strongly positively coregulated genes with the TSET core (average coregulation values > 0.8) include multiple endomembrane proteins associated with exocytosis: Phatr3_J43900, an ER membrane protein complex subunit 1; Phatr3_J10209, coatomer subunit gamma; Phatr3_J24186, exportin-1; Phatr3_J47327, exocyst complex component 7; and Phatr3_J49389, trafficking protein particle complex subunit 11. These may suggest possible roles for Phaeodactylum TSET in secretory processes. Intriguingly, the most positively correlated protein with the TSET core is a *P. tricornutum* uncharacterized protein with homology (at least within the C2-lipid binding domain) to another jotnarlog protein AGD12, a member of the ArfGAP_C2 protein subfamily [44], which acts within the Golgi body in plants and mediates gravitropism [45].

Further KEGG analysis of genes associated with core TSET subunits (r > 0.5, 2671 genes, 22% of total genes) confirms that the TSET complex may be associated with vesicular transport (Fig. 3). Indeed, the “Exosome”, “GTP-binding proteins” and “Membrane trafficking” categories ranked in the top 5 most complete functional categories (Fig. 3), with respectively 69% (106 coregulated /153 total genes in the “Exosome” category), 68% (13/19 genes) and 68% (181/266 genes) of completeness. Among the “GTP-binding proteins”, several genes are referenced to be directly involved in intracellular protein transport or to be part of the Ras superfamily (Phatr3_J43251, ARF1; Phatr3_J8659, ARL1; Phatr3_J54420, SAR1A; Phatr3_J13157; Phatr3_J45953). The “Photosynthesis proteins” category was ranked as the most complete category with 13 coregulated genes out of 15 (Fig. 3), which encodes proteins from the photosystems, the electron transport chain and the light-harvesting complexes. Based on previous studies about TSET functions, it is unlikely that this complex is directly linked to photosynthesis, but it may be involved in chloroplast biogenesis during cell division. Several metabolic pathways also have a strong association with the core TSET (data not shown), such as the “Signal Transduction” metabolic pathway (207 coregulated genes, 61% complete, 5^th^ rank) and the “Transport and Catabolism” metabolic pathway (175 coregulated genes, 60% complete, 6^th^ rank), the latter containing genes involved in endocytosis (34 genes), lysosomal activity (20 genes) and autophagy processes. These data suggest that in *P. tricornutum* the TSET complex is linked to key cellular processes such as vesicular trafficking, with possible implications in endo/exocytosis and cell division as demonstrated in *A. thaliana* and *Dictyostelium* [15,16].

**Figure 3.**
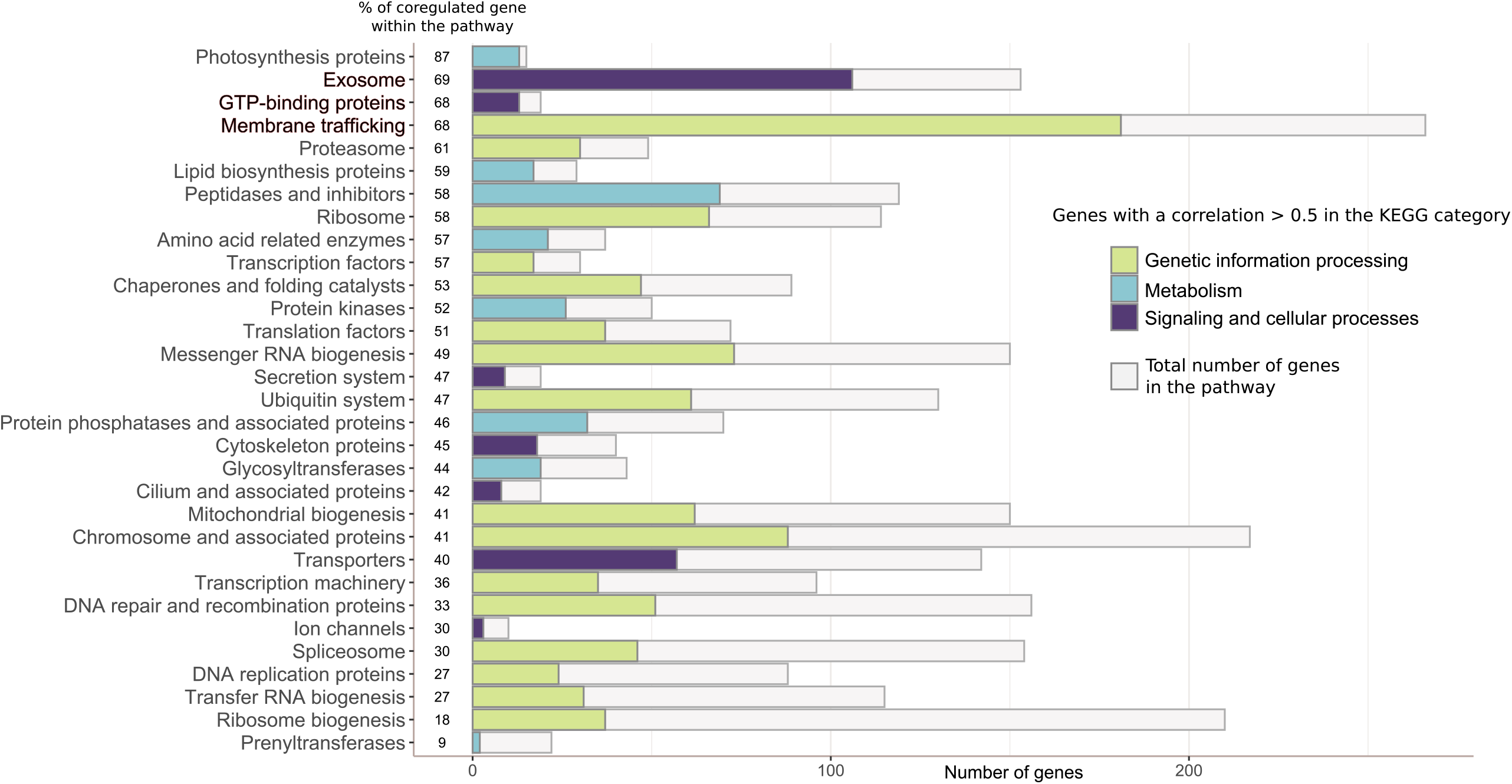
TSET expression in the diatom *Phaeodactylum tricornutum* is associated with endomembrane trafficking. Bar chart of enriched KEGG functions of genes found to be coregulated (r > 0.5) with four core TSET subunits (TCUP1, TPLATE, TSPOON1, TTRAY), assessed by Spearman correlation of published RNAseq and microarray data from the model diatom *P. tricornutum*. The numbers of coregulated genes in each category of the “Gene and proteins” classification of the KEGG Mapper Pathway are represented by colored bars, while grey bars represent the total number of genes in the corresponding category. Categories are sorted decreasingly from top to bottom based on their percentage of completeness (shown on the left of the bars). The “Enzymes” category (correlated = 579 genes, total = 1450 genes) and categories containing less than 5 genes are not considered. Raw supporting data are provided in Dataset S4.

### *Tara* Oceans data reveal the global presence and environmental significance of TSET

TSET components appear to be widely encoded in the genomes of at least some prominent marine algae, and there are indications from the co-regulation analysis of conserved cellular functions to those found in other eukaryotes. However, we wanted to understand whether TSET was playing a substantial role in real world settings. To assess this, we retrieved putative homologues of diatoms’ TSET proteins from metatranscriptome data gathered within the *Tara* Oceans expedition. This dataset encompasses both metagenomic and metatranscriptomic data over 153 globally distributed sites, separated by depth (surface, and Deep Chlorophyll Maximum), and cell size (from between 0.8 and 2000 µm diameter), allowing quantitative assessment of the roles of TSET in different vertical and size fractions of phytoplankton communities. Each sample within the *Tara* Oceans dataset is also associated with consistently measured environmental parameters (coordinate location, measured temperature, and measured or modelled concentrations of multiple nutrients including nitrogen, phosphorus, iron and silicon), allowing preliminary assessments of the physiological functions of TSET in natural communities.

The distribution of diatoms’ TSET subunits transcripts within the global ocean shows that TSET genes are expressed in the wild (Fig. 4). Over the 67 stations where putative TSET transcripts were detected, all of them presented the TCUP, TTRAY1 and TTRAY2 members, and 34 stations presented all the detected diatom subunits (Fig. 4). TSET components seem to be constitutively expressed in the wild diatom populations, the normalised abundances of the total putative TSET transcripts only showing a 5-fold variation magnitude (Fig. 4). Both the AP2 and TSET complexes are implicated in endocytosis from the cell surface. However, most models assessing heterotrophy only incorporate AP2 as the molecular mechanism for this process. In order to assess the relative rates of AP2 vs TSET, we compared the relative expression of the homologous medium subunits (TCUP vs AP2mu) within the *Tara* Oceans dataset. We observed that averaging across all size fractions and depth profiles, AP2mu is more highly expressed than is TCUP (Fig. S3). Nonetheless, the average TSET expression levels across all subunits, is comparable to that of enzymes in mitochondrial and plastid glycolysis [46] and so should be considered as a substantial facet to cellular function in diatoms in the oceans.

**Figure 4.**
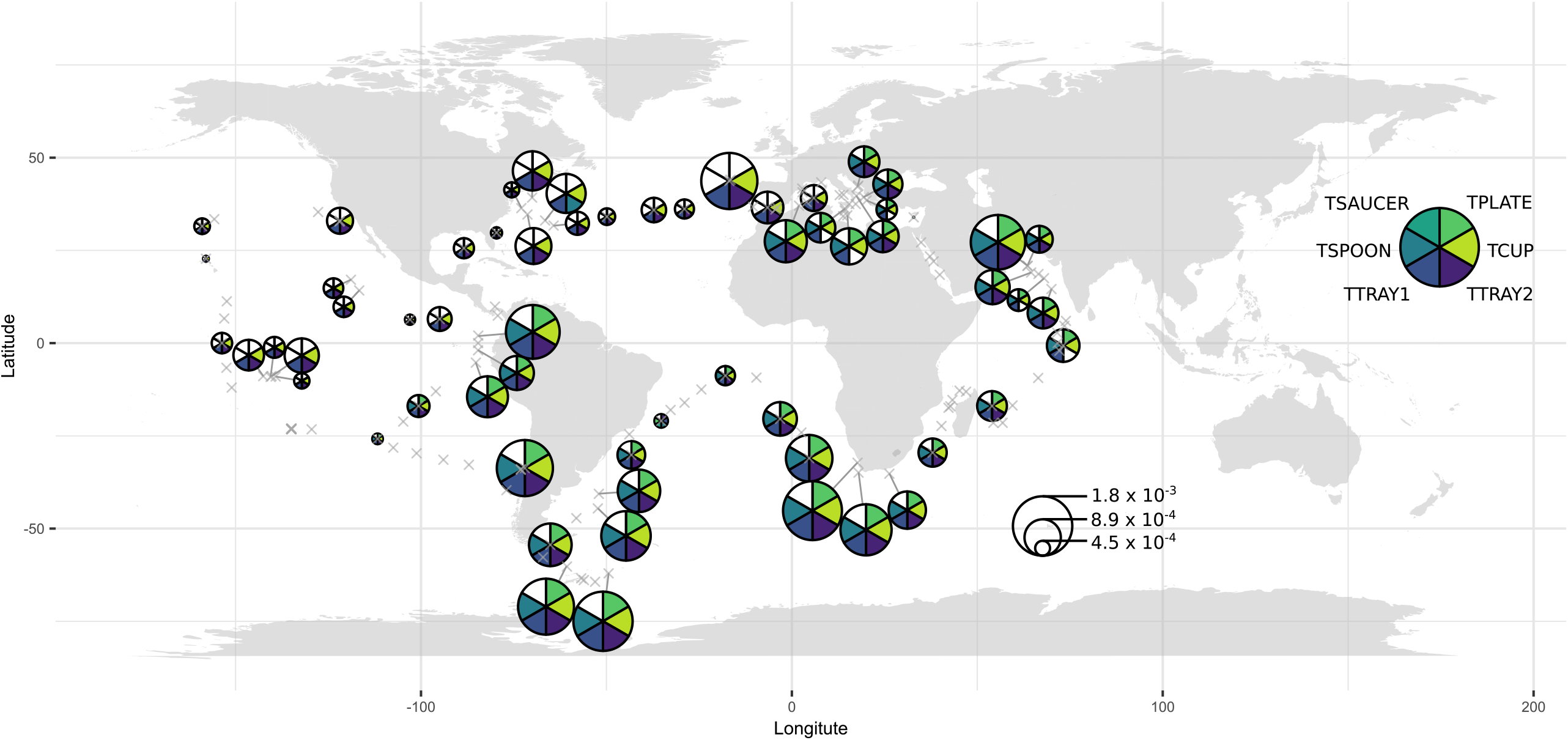
Map of *Tara* Oceans distributions of diatom TSET metatranscripts. Circles represent the mean between surface and DCM samples of the normalised total abundances of all putative TSET subunits transcripts found in each sampling station. The abundance of transcripts is expressed as the percentage of total transcript reads within the station, normalised by the total abundance of diatoms transcript reads within the same station. The presence or absence of the putative TSET subunits transcripts in a station is shown on the corresponding pie chart. Grey crosses indicate the location of all sampling stations. No occurrence of TSAUCER transcripts have been detected.

Visually, the TSET expression appears to be most abundantly expressed in stations from high latitude southern and coastal stations, and more variably highly expressed at northern stations, with lower normalised abundance in the tropical open ocean (Fig. 4). A global correlation analysis specifically projects negative correlations between TSET abundance and temperature, and positive correlations to oxicity and seawater density for multiple subunits (Fig. 5A). That said, a Principal Component Analysis (PCA) did not identify any one single environmental factor as having a predominant correlation with normalised TSET transcripts abundances (Fig. 5B). Overall, these data are consistent with diatom TSET being basally expressed, although this expression may be affected by a combination of measured and unmeasured environmental factors.

**Figure 5.**
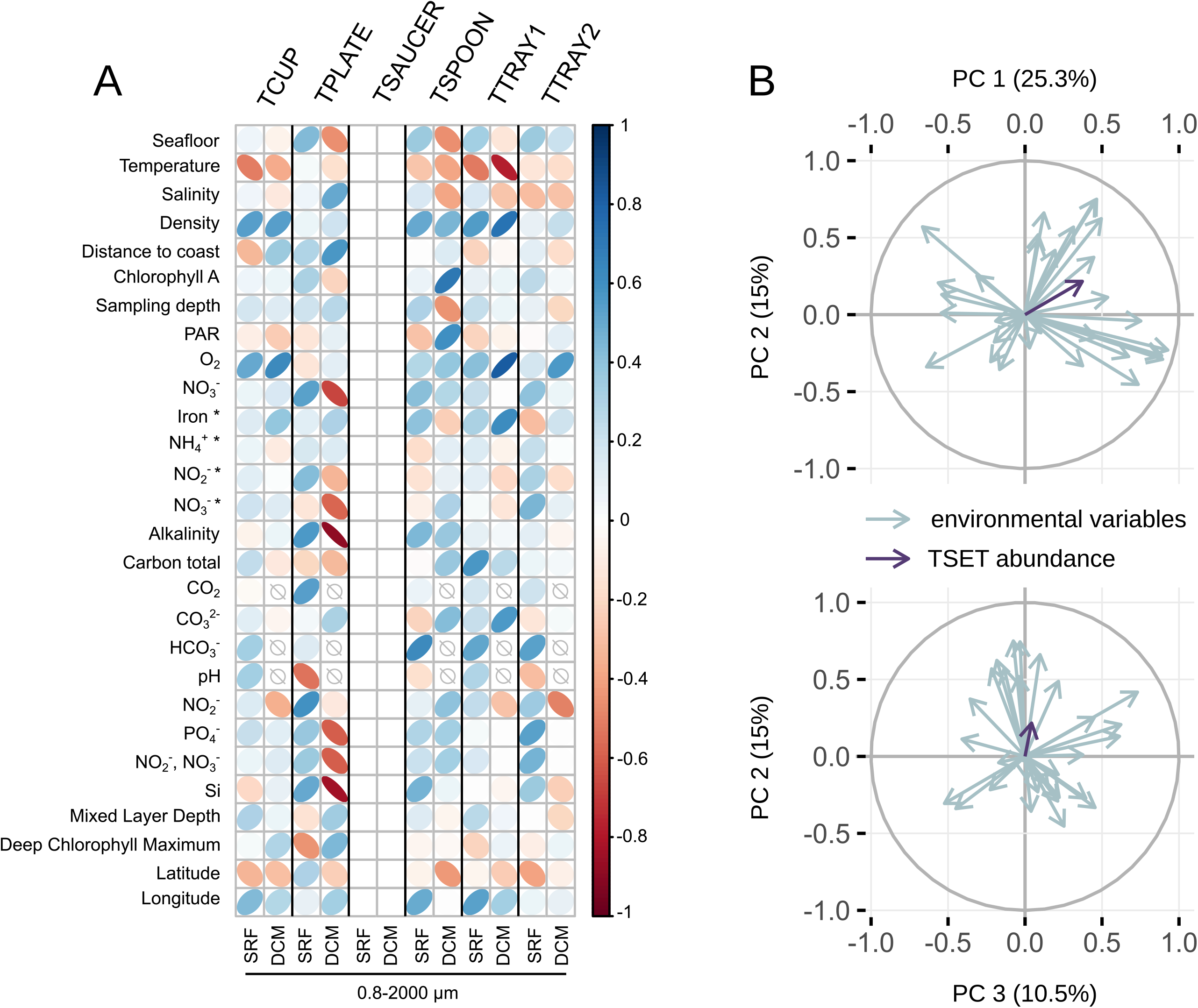
Correlations between the relative abundances of diatom TSET metatranscripts and *Tara* Oceans environmental variables A : Heatmap of the Spearman correlation coefficients between relative transcript abundances and environmental parameters. Normalised relative abundances for each sampling stations have been calculated for samples corresponding to the 0.8 to 2000 µm size fraction, collected at the surface (SRF) or at the Deep Chlorophyll Maximum (DCM). Shapes and colors of the ellipses correspond to the correlation coefficient value, empty signs show that no value is available for the corresponding environmental parameter. No occurrence of TSAUCER transcripts have been detected. B : Principal Component Analysis of *Tara* environmental conditions and TSET relative abundances. TSET abundances corresponds to the normalised total abundances of all putative TSET subunits metatranscripts found in each sampling station, depth and size fraction. Environmental parameters used for the PCA are listed in A, with the minimal and maximal O_2_ depths added. PC1 is positively associated with Nitrogen and Phosphorus nutrient concentrations, PC2 is positively associated with Carbon nutrient and pH, PC3 is associated with depth parameters (salinity, density, Photosynthetically Active Radiations).

## Conclusions

In this study, we harness systems-level and environmental data to explore the diversity and functions of a widely distributed but poorly understood membrane complex, TSET, in five environmentally important but evolutionarily heterogeneous protist and algal lineages.

Our results project much more widespread occurrence of TSET complex proteins in some of these groups than previously identified purely from small numbers of model species genomes. This seems particularly to be true for the green algae and stramenopiles, which both play substantial roles in marine ecosystems, as well as being of agricultural and medical significance in the case of stramenopiles. The gene co-regulation analysis strongly implicates TSET having a role within the endomembrane system of the model diatom *P*.*tricornutum*, despite our inability to identify the TSAUCER subunit in this organism, and many stramenopiles. From *Tara* Oceans data, we demonstrate that putative TSET genes are transcribed in wild population of diatoms, and that complete TSET (except TSAUCER) can be virtually found in many stations across the world. No specific contributions of environmental parameters have been detected in transcript abundances, suggesting that TSET expression may either be constant, or under complex regulation inside the diatom cell.

There are clear limitations to the study to declare. Firstly, because our dataset was designed to maximize taxonomic inclusion, there is a clear danger of false negatives in our results despite our attempts this mitigate with multiple search strategies. This could be due to sequencing depth, combined with low expression of TSET subunits in the conditions under which the organisms were cultured. Still, the overall paucity of identified subunits in the red algae, cryptophytes, and haptophytes is not consistent with a major role for TSET in those taxa, with the caveat of the speculated TSPOON monomeric protein in haptophytes. Nonetheless, the highly sporadic distribution of TSAUCER in stramenopiles (with individual datasets possessing validated orthologues and other datasets from closely related datasets lacking them), and the evidence that the other complex subunits are co-regulated and expressed in the wild, all suggest that there may be more to this story. Either a highly divergent TSAUCER subunit is present and escaping detection by our informatic methods, or some analogous protein has been recruited. Secondly as a separate limitation to discuss, though we have tried to be conservative, limiting ourselves to ‘involvement in endomembrane pathways’, any inferences about the cellular role of TSET in *P. tricornutum* are speculative at best. This is due to the indirect nature of gene co-regulation networks and the unique biology of plants which is almost the only available molecular cell biologically characterized system against which to compare. The potential functions of TSET in *Phaeodactylum* or indeed other algae, will be best investigated by reverse genetic strategies, which are now routine for this lineage. These may include the experimental localisation of GFP-linked constructs across the duration of the cell cycle, or generation and phenotypic analysis (e.g., growth or metabolomic analysis of osmotrophic capacities) of CRISPR knockout lines. Similarly, either IP-pulldowns or sub-cellular proteomics could address the question of whether a TSAUCER orthologue or analogue exists, in addition to identifying a wealth of additional interaction data, as was recently done for *A. thaliana* [47].

More broadly, our data reinforce two major themes. Firstly is the importance of using phylogenetically diverse references, including both cultured and uncultured microbial species, for understanding the significance of broadly conserved eukaryotic cellular processes. The second is that jörtnarlog genes, at least in the case of TSET, can be quite widespread in groups of environmentally important groups of microbial eukaryotes, and that these genes are being expressed in wild populations. Study of classical model species within the opisthokonts provided deep comprehension of a set of cellular mechanisms. And yet, as with all eukaryotic lineages opisthokonts have their own uniquely evolved cell biology, which is not always applicable to the rest of the eukaryotes. The heavy reliance on opisthokont models have created paradoxical situations where widespread genes within eukaryotes (jötnarlogs) can still have unresolved function. Development of model organisms from other parts of the eukaryotic tree, led by studies in embryophytes, and a few key parasites have been instrumental to bringing this bias to light. However, because plants have unique multicellular biology, they are often unfairly discounted as general eukaryotic models. Parasites such as Toxoplasma, Trypanosoma, and to a lesser extent Giardia, Plasmodium, Trichomonas and Entamoeba can be powerful comparators to better understand eukaryotic cell biology, with a rich set of genomic, cell biological, and system biology tools and data available. However, because such organisms are often highly reduced, diverged, and/or specialized, the homology of their cellular systems has been cryptic and underused in developing general cell biological models. Where there is clear concurrence with opisthokont cell biology, it is integrated,. Where there is discordance, the situation becomes more complicated. We are advocating for an even broader perspective: integrating algae, protists and phylogenetically diverse eukaryotic organisms in cell biology research [11] could provide several benefits to this field. These may include access to a wide range of behavioural and trophic strategies, or a better proxy to understand molecular coevolution. Moreover, *Tara* Oceans and other similar sampling campaigns provide not only large but repeatable data, giving access to huge amounts of quantitative expression data alongside clearly and consistently measured environmental variables. We believe that these ecologically important microscopic organisms, and their uncultured environmental relatives, combined with this wealth of underexplored proteins (jötnarlogs) in membrane trafficking and other cellular systems may hold particular promise in revealing fundamental processes in evolutionary cell biology.

Dataset S1: Composition of the multispecies reference library and taxonomic divisions used for TSET comparative genomic analysis. Data are available at https://osf.io/ykxes/.

Dataset S2: List and sequences of query TSET protein used for homologue retrieval with AMOEBAE.

Dataset S3: List of TSET homologs, alignments and curated tree topologies, and Ref_seqs_1_manual_predictions.csv used for the AMOEBA searches.

Dataset S4: Coregulation coefficients of six recovered TSET subunit genes -Phatr3_J43047 (TCUP1), Phatr3_J43761 (TCUP2), Phatr3_J54511 (TPLATE), Phatr3_J54718 (TSPOON1), Phatr3_J14536 (TSPOON2), and Phatr3_J46356 (TTRAY) – and all other genes in the Phaeodactylum tricornutum genome version 3 annotation, assessed by Spearman correlation analysis of merged RNAseq (PhaeoNet) and microarray (DiatomPortal) data.

Sheet 1 (“Phat3 genome”) provides an overview of the entire genome and its associated transcriptional dynamics.

Columns A to C provide each gene, its ID and full length peptide sequence. The first six rows correspond to inferred TSET homologues.

Columns D to M provide information concerning the expression trends associated with each gene: its occurrence in DiatomPortal and PhaeoNet coexpressed gene modules; and its calculated Spearman coregulation coefficient to each inferred TSET gene, all six TSET genes (“TSET average”) or the average coregulation coefficient calculated purely for TCUP1, TPLATE, TSPOON1, and TTRAY (“Core average”).

Columns N to R provide information concerning the inferred localisation of the protein encoded by each model, considering the consensus of ASAFind (realised with SignalP 3.0), HECTAR, MitoFates (with threshold value 0.35) and WolfPSort (consensus animal, plant and animal annotations).

Columns S to Z provide information concerning the inferred function of the encoded protein, considering KEGG annotation (based on BLASTKOALA, GhostKOALA, KofamKOALA), PFAM and predicted biological process.

Columns AA to AE provide associated genomic contexts, considering chromosomal localisation, epigenetic marks (histone methylation and acetylation, and DNA methylation), and presence of inferred alternative splice-forms from mapped RNAseq data.

Sheet 2 (“Core enrichments”) provides a chi-squared pivot table of PhaeoNet RNAseq WGCNA merged modules, consensus localisation predictions, and consensus KEGG annotations enriched in genes with moderate (r> 0.5) or strong (r> 0.7) positive coregulation to the four core TSET subunits (TCUP1, TPLATE, TSPOON1, TTRAY), as defined above. P-values are signed so that occurrences that are less frequently observed than expected are negative, and occurrences that are more frequently observed than expected are positive.

Dataset S5: Table of AMOEBA search results for TSET subunits, sorted by taxonomy. Only genomes/transcriptomes with two or more AP1 subunits detected are presented.

Dataset S6: Tabulated Tara Oceans TSET homologues and relative abundances in different stations, depths and size fractions.

## Supporting information

Compressed files for Dataset S3

---

**Figure S1:**
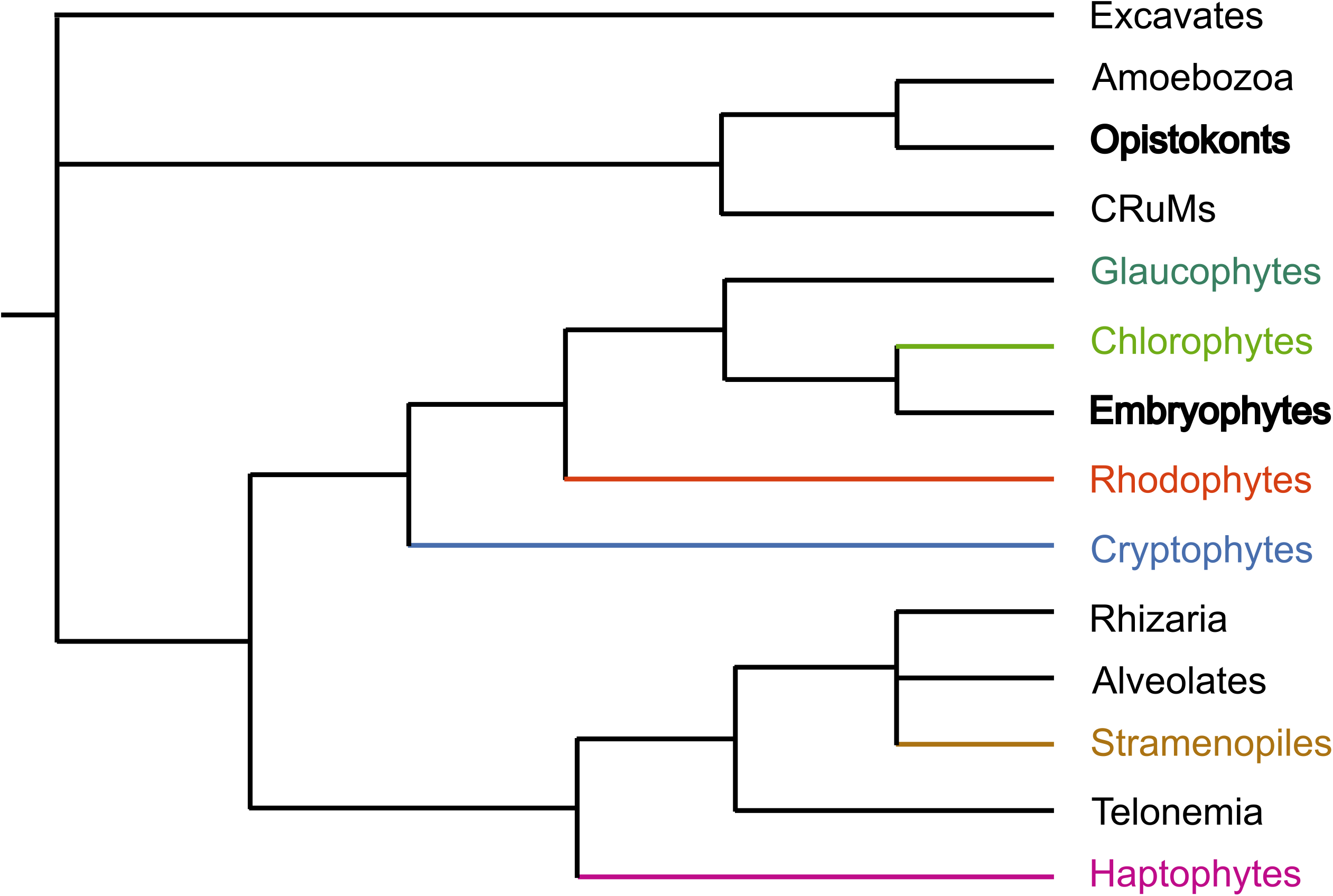
Schematic tree of Eukaryotes. Taxa containing classical model organisms in cell biology are in bold, coloured taxa are the ones on which this study focuses. Branch lengths are arbitrary, and exact tree topology is still under debate.

**Figure S2:**
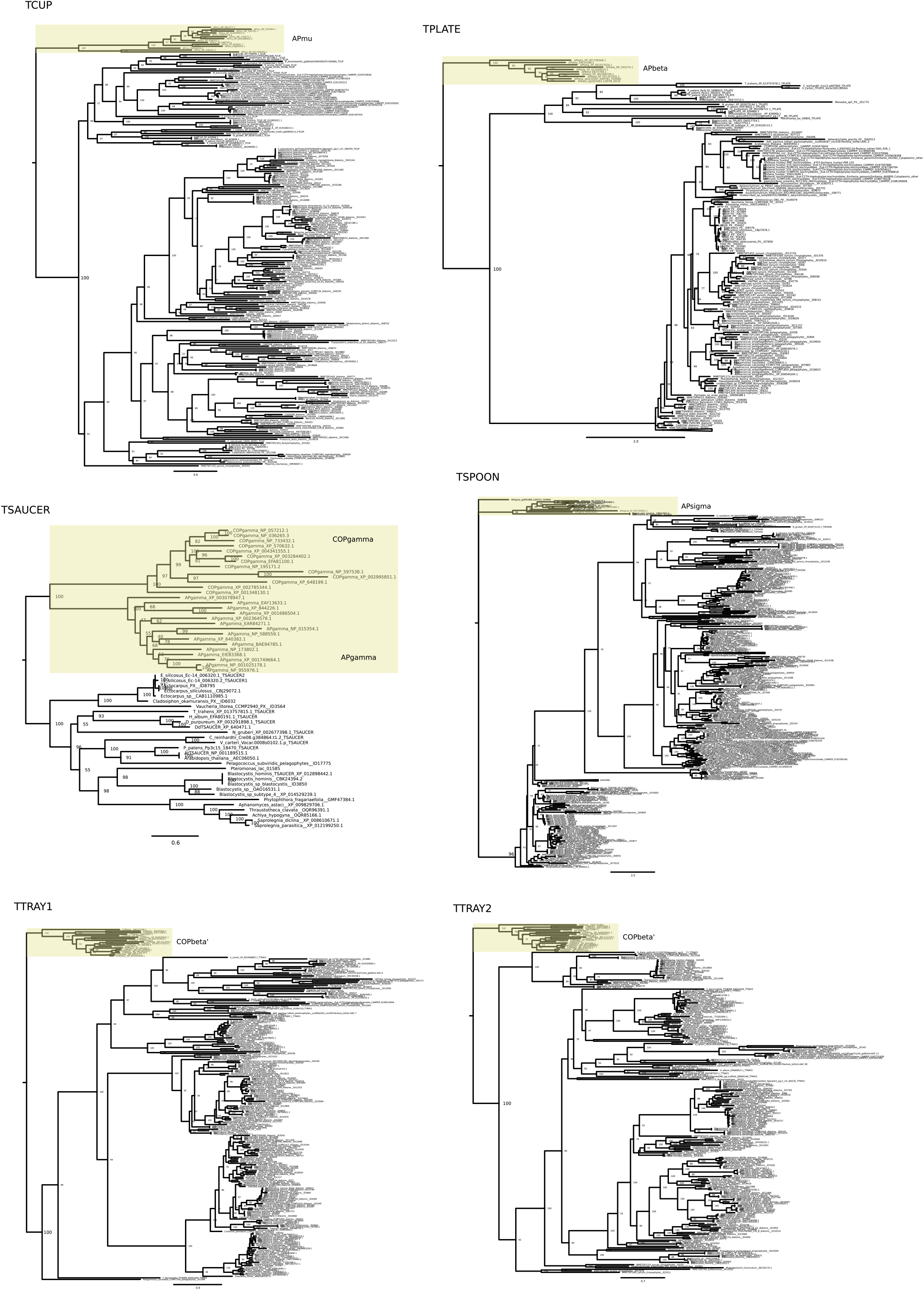
Consensus trees of stramenopile and haptophyte TSET subunits. Phylogenies were computed with IQ-TREE2 v2.2.6. The VT+R7 model (best model assessed by ModelFinder for TCUP) was used for each tree. Branch support are ultrafast bootstrap approximation (1000 replicates). Outgroups correspond to homologous AP or COP subunits

**Figure S3:**
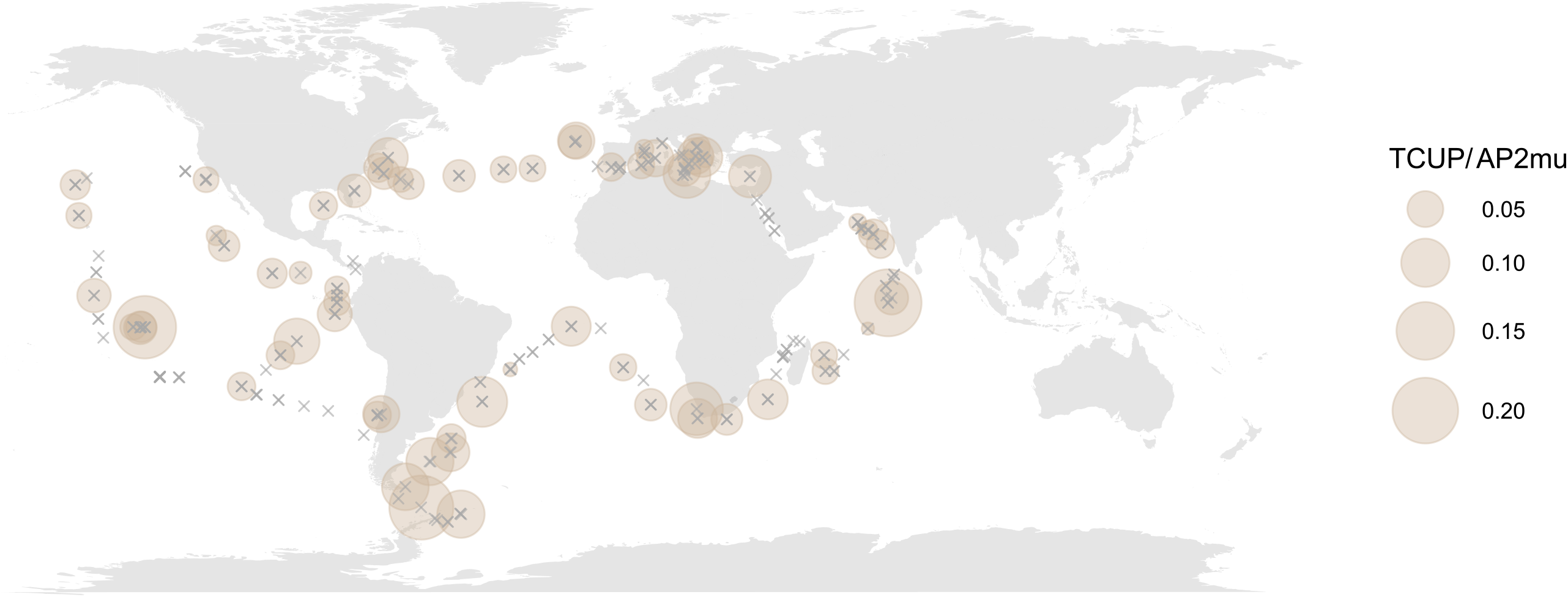
Map of *Tara* Oceans distributions of stramenopile TCUP vs AP2mu metatranscripts. Circles represent the ratio between the mean total abundances of all putative TCUP and APmu subunits transcripts found in each sampling station. The abundance of transcripts is expressed as the percentage of total transcript reads within the station. Grey crosses indicate the location of all sampling stations.

